# SBP-SITA: A sequence-based prediction tool for S-itaconation

**DOI:** 10.1101/2021.12.13.472522

**Authors:** Laizhi Zhang, Xuanwen Wang, Lin Zhang, Yanzheng Meng, Ziyu Wang, Yu Chen, Lei Li

## Abstract

As a recently-reported post-translational modification, S-itaconation plays an important role in inflammation suppression. In order to understand its regulatory mechanism in many life activities, the essential step is the recognition of S-itaconation. However, it is difficult to identify S-itaconation in the proteome for the high cost, which limits further investigation. In this study, we constructed an ensemble algorithm based on Soft Voting Classifier. The area under the ROC curve (AUC) value 0.73 for ensemble model. Accordingly, we constructed the on-line prediction tool dubbed SBP-SITA for easily identifying Cystine sites. SBP-SITA is available at http://www.bioinfogo.org/sbp-sita.

## 1 Introduction

As one of the 20 amino acid composing protein, cysteine has high reactivity for its thiol side chain. Some cysteine residues in protein could covalently bind to endogenous molecules like succinate or itaconate, forming post-translation modification (PTM). Itaconate is a metabolite molecule, which was first observed to be present in mammalian cells during macrophage activation in 2011 (Strelko et al., 2011). Six years later, it was found that itaconate could also modify cysteine residue in proteins, resulting in unreversible PTM S-itaconation (Mills et al., 2018). S-itaconation regulate a series of immune process including decreasing antigen presentation, anti-inflammation and decreasing glycolysis (O’Neill and Artyomov, 2019). Recently, its unusual role in tumor development was also been revealed (Weiss et al., 2018).

Because of the functional significance of S-itaconation, the identification of new S-itaconatoin sites in proteins and proteome is much more important. However, antibody that could capture S-itaconate proteins or peptides are still lacking. Although some chemical proteomics techniques were developed to identify S-itaconate protein from cell lysates, they are expensive and time-consuming (Qin et al., 2019; Qin et al., 2020). By contrast, combing big data and advanced algorithms, some bioinformatics method can provide an alternative strategy by predicting S-itaconation. Some bioinformatics prediction tools have been developed for certain types of cysteine PTM like DeepCSO for S-sulphenylation, GPS-Palm for S-palmitoylation and so on (Lyu et al., 2020; Ning et al., 2020). These bioinformatics predictors become complement approaches for wet experiment, promoting the investigation of different kinds of cysteine PTM. Nevertheless, no prediction model is available for the S-itaconation.

In this study, we developed a machine-learning prediction tools named SPS-SITA. We firstly collect the identified 997 S-itaconation sites in RAW 264.7 cell lines to make dataset and then developed an Ensemble predictor by Soft Voting based on six models. It achieves AUC 0.73. We also provided the online SBP-SITA via https://bioinfogo.org/sbp-sita

## 2 Materials and Methods

### 2.1 Data Collection and Preprocessing

In this study, we collected the experimentally identified S-itaconation protein sequences from literature (Qin *et al*., 2020). To construct positive data of modeling, a sequence window was applied to intercept the protein sequences. After this process, we obtained an amino acid sequence of length 27, with the modified cysteine site in the center (position 14), containing 13 amino acids on each side. If there were less than fifteen amino acids on one side, we used “*” to replace the lacking residues. Afterwards, negative data are generated using a similar methodology, where the unmodified cysteine was in the center of the sequence windows (Wang et al., 2020). A data set containing 1422 positive sites and 17414 negative sites was established. Second, to eliminate the homology bias, CD-HIT tool was employed with 40% identity. Redundant samples in the data sets were removed that that none of the sequences had a larger than 40% pairwise identity in both positive and negative sets. Thirdly, to prevent errors caused by data imbalance, we performed a down-sampling for the data. Specifically, after CD-HIT process we obtained 995 positive peptides and 5140 negative peptides. In order to equalize the positive and negative sample sizes, we randomly selected the same number of negative sequences as the positive samples using down-sampling, i.e., 995 negative samples were randomly chosen.

In the end, we obtained 1990 sequences from mice for modeling. To effectively assess the performance of models, 10-fold cross validations and independent testing were employed on the mouse data (Chen et al., 2018a). We randomly selected 398 samples from 1990 protein fragments of mouse data for independent testing. Then the remaining 1592 sequences were randomly divided into deciles. One set of data was used for model validation, while the remaining nine sets of data served as the training set (Figure 1A).

**Figure 1.**
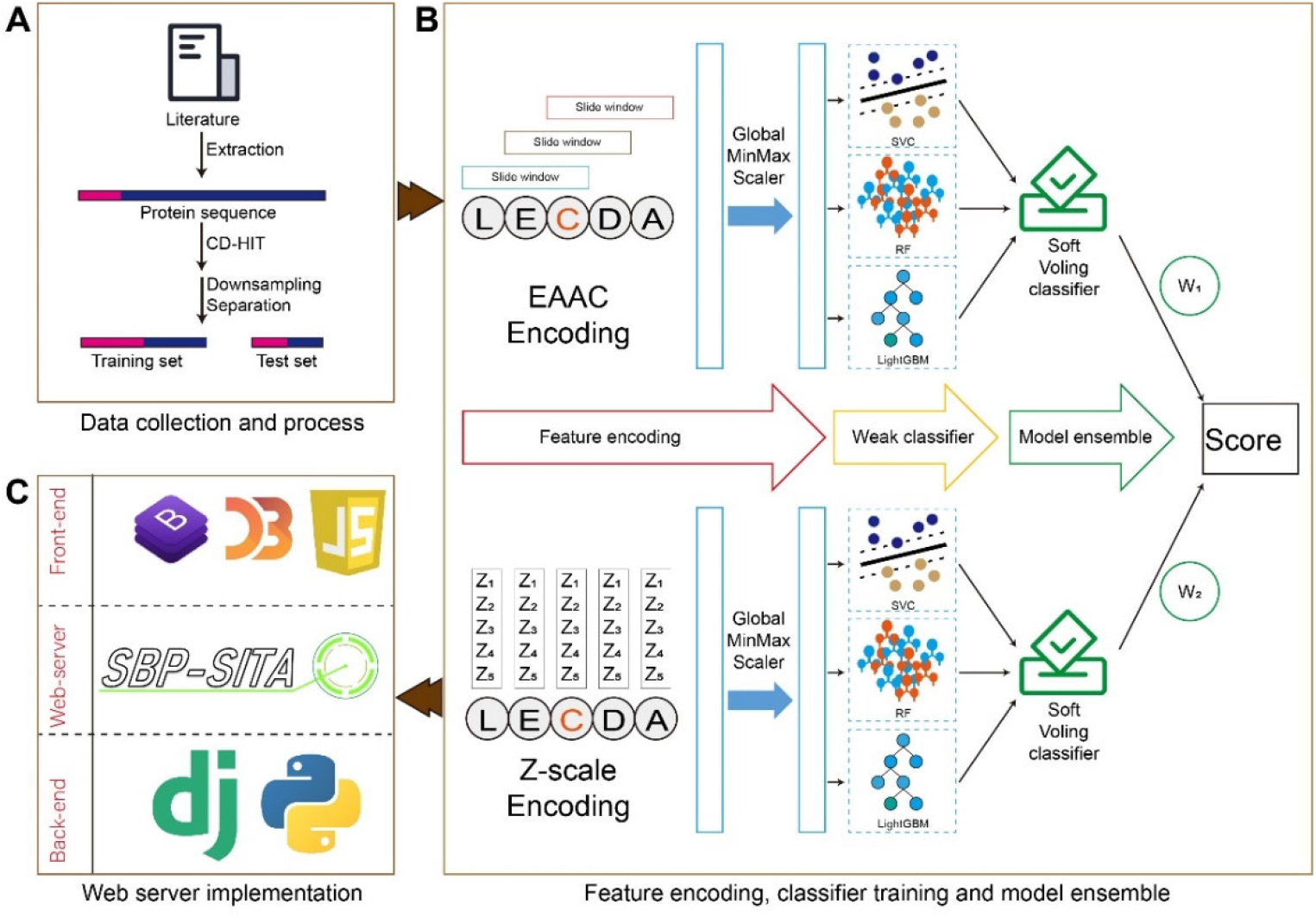
(A) From the scientific literature we collected S-itaconation data set. After redundancy clearance and downsampling, only 1994 sequences were remained as data sets. (B) Six models were trained and integrated as an Ensemble model by Soft Voting. (C) The web-server implementation of SBP-SITA.

### 2.2 Feature Investigation

To check the quality of the training data, we visualized the features of the sequences to facilitate our inspection via Two Sample Logo web server (Vacic et al., 2006). It calculated the statistical significance of the position specific residue frequencies between the positive and negative sequences in training sets, showing the significantly enriched or depleted residues in the flanking sequence of center cysteine. To be more specific, in the output graph, the upper part shows the set of characters which are enriched (overrepresented) in the positive set. The lower part displays the set of characters which are depleted (underrepresented) in the positive set while the middle part shows the consensus characters. We also used this with model discission ratio the importance of each position in sequence through sequence encoding. (Fig 3A)

### 2.3 Sequence Encoding

In order to facilitate computer processing of amino acid sequences, we converted the sequence information into a format that can be processed by programs. Therefore, the amino acid sequences represented by letters is converted into quantification vectors. We used the following four methods to encode the sequences in order to preserve the positional information as much as possible and to facilitate the extraction of peptide features.

#### 2.3.1 Binary Encoding (BE)

Binary encoding (BE) uses 0, 1 for representing the twenty amino acids. For each amino acid letter on the peptide, there is a 20-dimensional vector of zeros and one, i.e., only one position is 1 and the remaining 19 positions are set as zero (Chen et al., 2018b). For example, (0100000000000000000000) is used to represent C, and (00000000000000000001) is used to represent Y, uniquely. When we choose the input sequence length of 51 amino acids, after BE process we got the vector with the shape of (51, 20).

#### 2.3.2 Enhanced Amino Acid Composition (EAAC)

EAAC was developed based on Amino Acid Composition (AAC) encoding method, which counts the frequency of distribution of each amino acid on the sequence (Chen *et al*., 2018a). Unlike BE, AAC cannot cover the position information of amino acids. Therefore, EAAC solves this problem by introducing a sliding window of fixed length over the full length of the peptide, and then based on the statistical method of AAC. For instance, when a fragment of 31 amino acids in length was input, by EAAC encoding (with a sliding size of default 6) we end up with a vector of shape (26, 20).

#### 2.3.3 Composition of k-spaced Amino Acid Pairs (CKSAAP)

The CKSAAP (Composition of k-spaced Amino Acid Pairs) method builds a mathematical model for feature vector extraction using the proportion of k-spaced residue pairs in a fragment of a protein sequence (Chen et al., 2020). Taking a peptide of length 51 as an example, when k=0, we can extract 50 residue pairs with 0 amino acids in each residue pair spaced in the sequence. And so on, when we choose k to be 1, 49 residue pairs can be extracted, so when the number of spaced amino acids is k, we can extract:

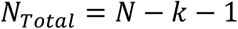

Since the number of basic amino acids is 20, the number of residue pairs that can be formed is 20 × 20 = 400. We calculate the probability that these pairs of residues occur in a given protein sequence, resulting in a 400-dimensional feature vector. It can be defined as:

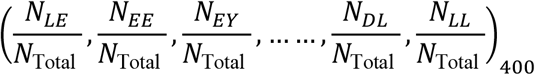

We end up with a vector of the shape (400,).

#### 2.3.4 Z-SCALE (ZSCALE)

Z-Scale method was developed by Sandberg et al. in 1998 which embeds the physicochemical properties of amino acids into vectors (Sandberg et al., 1998). The ZSCALE descriptor can be applied to encode peptides of equal length. When we input a peptide of 51 amino acids in length for encoding, we end up with a vector of shape (51, 5) through the Z-S descriptor (Table S1).

### 2.4 Model Developing

After the protein fragments were encoded, we used the following four different algorithms to fit the training set and then to predict the input sequences (Figure 2B).

**Figure 2.**
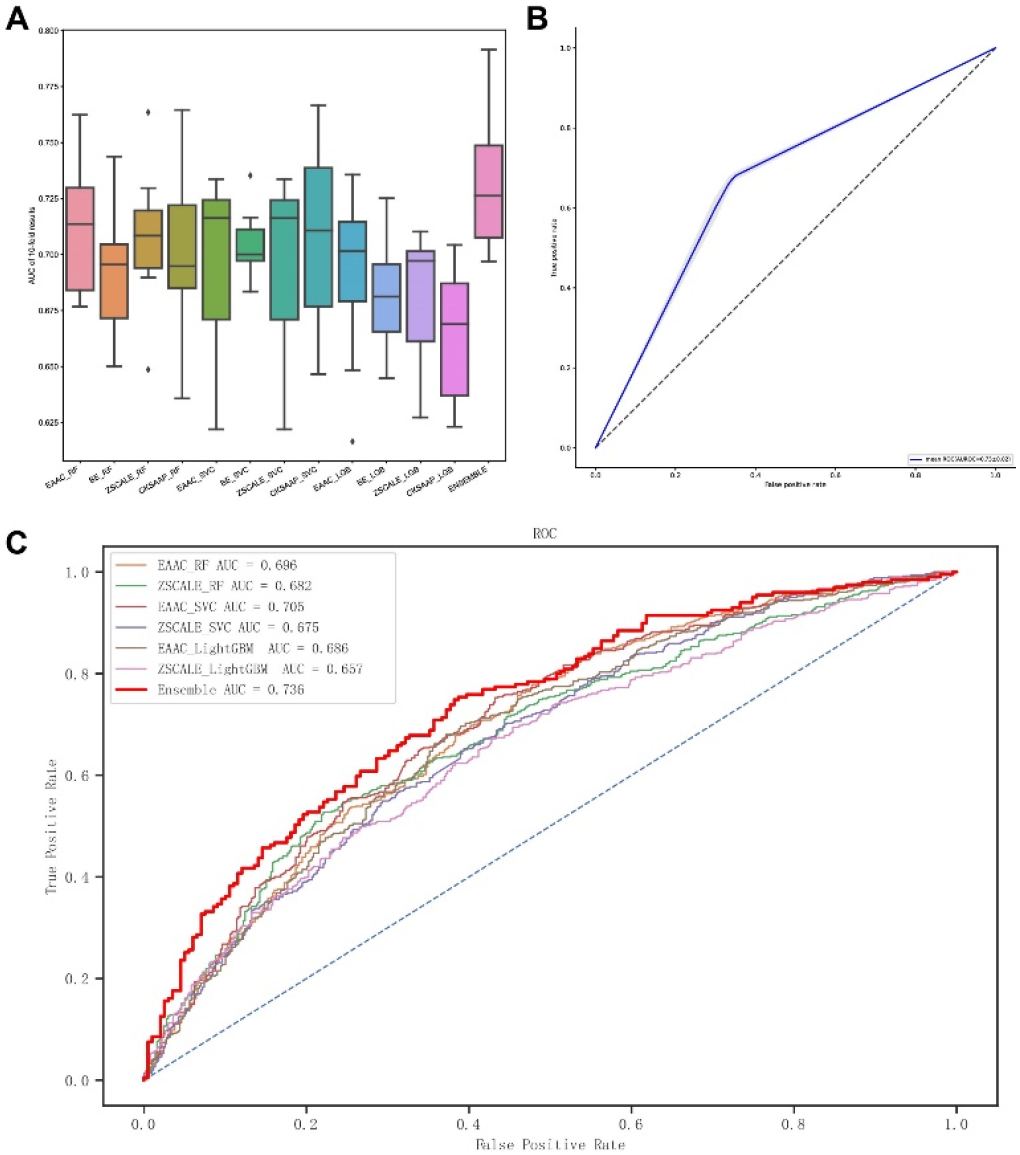
(A) The average AUC of each models for 10-fold cross-validation. (B) The average AUC of Ensemble in independent test. (C) The comparison of Ensemble and six models.

#### 2.4.1 Support Vector Classifier with Different Encoding Features

Support vector classifier (SVC) is a two-class classification model. Its basic model is a linear classifier defined by maximizing the interval on the feature space; support vector machines also include kernel tricks, which make it a substantially nonlinear classifier (Weng et al., 2017). The learning strategy of a support vector classifier is interval maximization, which can be formalized as a problem of solving convex quadratic programming, which is also equivalent to the minimization of a regularized hinge loss function. The learning algorithm for it is the optimization algorithm for solving convex quadratic programming. It can solve the problem of high-dimensional, nonlinear machine learning with small samples and improve the generalization performance, and also avoid the complex problems of neural network structures.

#### 2.4.2 Random Forest Classifier with Different Encoding Features

The Random Forest (RF) algorithm contains several decision trees that remain invariant under scaling of eigenvalues and various transformations, with the output class determined by the output class pattern of individual trees (Huang et al., 2018). Each tree depends on the values of a random vector with the same distribution of all trees in the forest. As with the SVM, we used the results of the four encodings (BE, EAAC, CKSAAP, Z-SCALE) as input to the RF. This classifier is based on the Python “sklearn” module developed for prediction. And we use the “GridSearchCV” method for optimal hyperparameter selection. Finally, we obtain a decision tree and thus the importance of each feature that can be interpreted.

#### 2.4.3 LightGBM with Different Encoding Features

LightGBM is a gradient boosting framework that uses tree-based learning algorithms. The boosting tree uses an additive model with a forward stepwise algorithm to implement the optimization process of learning. When the loss function is squared error loss function and exponential loss function, each step of optimization is simple. However, for general loss functions, it is often not so easy to optimize each step. To address this problem, Freidman proposed the gradient boosting algorithm. Gradient Boosting is a large class of algorithms in Boosting, which borrows its idea from gradient descent, and its basic principle is adding weak classifiers based on the negative gradient information of the loss function for the current model and then combine the trained weak classifiers into the existing model in an accumulative form (Friedman, 2001).

#### 2.4.4 Model ensemble with Soft Voting Classifiers

Voting Classifier is an estimator that combines models representing different classification algorithms associated with individual weights for confidence (Tasci et al., 2021). Voting classifier takes majority voting based on weights applied to the class or class probabilities and assigns a class label to a record based on majority vote. The ensemble classifier prediction can be mathematically represented as the following:

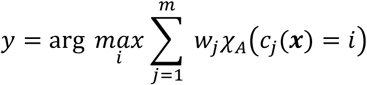

### 2.5 Model Implement and Classifier Performance Evaluation

After determining the algorithmic framework of models, we performed ten-fold cross-validations on the training data to select the optimal hyperparameters using grid search, while evaluating the stability of the model. To evaluate the generalization ability of the models, we predicted and scored the tuned completed models on the independent test set. The following five performance evolution indexes are reported. For ten-fold cross-validation, the mean and standard deviation of each index are described:

ACC:

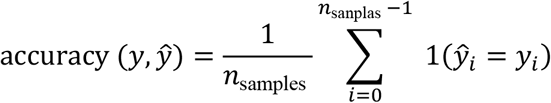

AUC:

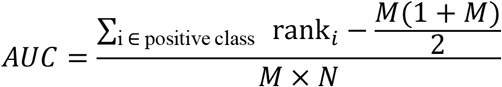

SP:

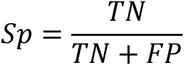

SN:

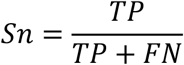

F1-Score:

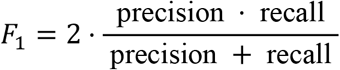

## 3 Results

### 3.1 EAAC_RF Model Performed Favorably to Other Single Models

Many prediction models for predicting S-itaconation sites were trained based on three ML algorithms combined with four features encoded from peptide sequences in this study. For each model, their performances of these algorithms in terms of several measures including AUC, ACC, SP, SN and F1-Score for 10-fold cross-validation were recorded (Table 1). To find the best model, we compare their average AUC values in 10-fold cross validation. Among these models, EAAC_RF shows superior performance than other single models, which had the average AUC values of 0.71 (Figure 2A). For SVC and LightGBM, EAAC coding were also better than other encoding method with 0.70 and 0.69 average AUC values, respectively. ZSCALE performed also well, for the potential reason that Zscale containing chemical information of each amino acids. BE performed significantly lower than the other encoding methods, indicating that it was not applicable to this experiment.

**Table 1.** 10-fold Validation Performance.

| method | AUC | ACC | SP | SN | F1-score |
| --- | --- | --- | --- | --- | --- |
| EAAC_RF | 0.712±0.030 | 0.657±0.027 | 0.660±0.060 | 0.652±0.061 | 0.654±0.038 |
| BE_RF | 0.693±0.030 | 0.638±0.029 | 0.629±0.083 | 0.652±0.052 | 0.643±0.030 |
| ZSCALE_RF | 0.707±0.029 | 0.641±0.020 | 0.622±0.059 | 0.660±0.053 | 0.647±0.029 |
| CKSAAP_RF | 0.699±0.043 | 0.639±0.038 | 0.675±0.056 | 0.602±0.058 | 0.624±0.046 |
| EAAC_SVC | 0.696±0.038 | 0.646±0.029 | 0.655±0.044 | 0.636±0.054 | 0.642±0.036 |

Table 1 10-fold Validation Performance
|  |  |  |  |  |  |
| --- | --- | --- | --- | --- | --- |
| BE_SVC | 0.703±0.015 | 0.647±0.019 | 0.660±0.036 | 0.633±0.057 | 0.640±0.035 |
| ZSCALE_SVC | 0.696±0.037 | 0.644±0.032 | 0.658±0.043 | 0.630±0.059 | 0.638±0.041 |
| CKSAAP_SVC | 0.708±0.038 | 0.646±0.036 | 0.697±0.042 | 0.594±0.059 | 0.626±0.049 |
| EAAC_LightGBM | 0.691±0.036 | 0.644±0.037 | 0.633±0.057 | 0.654±0.047 | 0.647±0.036 |
| BE_LightGBM | 0.682±0.023 | 0.633±0.023 | 0.624±0.049 | 0.642±0.049 | 0.635±0.035 |
| ZSCALE_LightGBM | 0.680±0.031 | 0.628±0.036 | 0.634±0.078 | 0.620±0.051 | 0.624±0.034 |
| CKSAAP_LightGBM | 0.665±0.029 | 0.625±0.022 | 0.616±0.045 | 0.633±0.061 | 0.627±0.037 |
| Ensemble | 0.733±0.030 | 0.672±0.046 | 0.676±0.064 | 0.667±0.073 | 0.669±0.054 |

### 3.2 Establishment of the Ensemble Model via Soft Voting

Due to the potential complementary effects in combining different classifiers to achieve better results, we investigated whether an integration of more classifiers would be more robust or perform better. We developed an Ensemble classifier by integrating six models (RF_EAAC_, SVC_EAAC_, LightGBM_EAAC_, RF_ZSCALE_, SVC_ZSCALE_ and LightGBM_ZSCALE_) via Soft Voting (Figure 1B). Ensemble model showed outstanding performance for both cross-validation and the independent test (Figure 2A, Figure 2B, Table 1, Table2). The AUC value was 0.73 in both 10-fold cross validation and test, which was better than six single models integrated in it. This model was chosen as the final model.

**Table 2.** Independent Test Performance.

| method | AUC | ACC | SP | SN | F1-score |
| --- | --- | --- | --- | --- | --- |
| EAAC_RF | 0.716017 | 0.663317 | 0.658291 | 0.668342 | 0.665 |
| BE_RF | 0.661069 | 0.615578 | 0.653266 | 0.577889 | 0.600522 |
| ZSCALE_RF | 0.695437 | 0.645729 | 0.638191 | 0.653266 | 0.648379 |
| CKSAAP_RF | 0.7043 | 0.660804 | 0.728643 | 0.592965 | 0.636119 |
| EAAC_SVC | 0.689212 | 0.638191 | 0.643216 | 0.633166 | 0.636364 |
| BE_SVC | 0.693947 | 0.640704 | 0.643216 | 0.638191 | 0.639798 |
| ZSCALE_SVC | 0.689326 | 0.638191 | 0.648241 | 0.628141 | 0.634518 |
| CKSAAP_SVC | 0.715373 | 0.663317 | 0.723618 | 0.603015 | 0.641711 |
| EAAC_LightGBM | 0.676751 | 0.635678 | 0.638191 | 0.633166 | 0.634761 |
| BE_LightGBM | 0.673948 | 0.640704 | 0.653266 | 0.628141 | 0.636132 |
| ZSCALE_LightGBM | 0.655691 | 0.628141 | 0.663317 | 0.592965 | 0.614583 |
| CKSAAP_LightGBM | 0.654403 | 0.623116 | 0.592965 | 0.653266 | 0.634146 |
| Ensemble | 0.751471 | 0.693467 | 0.693467 | 0.693467 | 0.693467 |

### 3.3 Feature Exploring of S-itaconation

Two Sample Logo was employed to show the differences of amino acid distribution in each position between positive data set and negative data set (Figure 3A). The larger fonts denote the amino acid tending to appear in this position with statistical significance. It was shown that the amino acids with polarities, especially Glycine (G) and Serine (S), and one nonpolar amino acid Proline (P) were enriched in the six amino acids around S-itaconation sites. We also counted the feature importance of BE encoding. For BE feature importance, position 13, 15, 17, 12 contributed mostly to S-itaconation, showing the consistent position location of Two sample logo (Figure 3B). To further explore it occurring in physicochemical information, Z-scale feature importance was also employed. It also showed similar trends in Two sample logo, indicating the amino acids near the Cystine may influence it via physicochemical effect.

**Figure 3.**
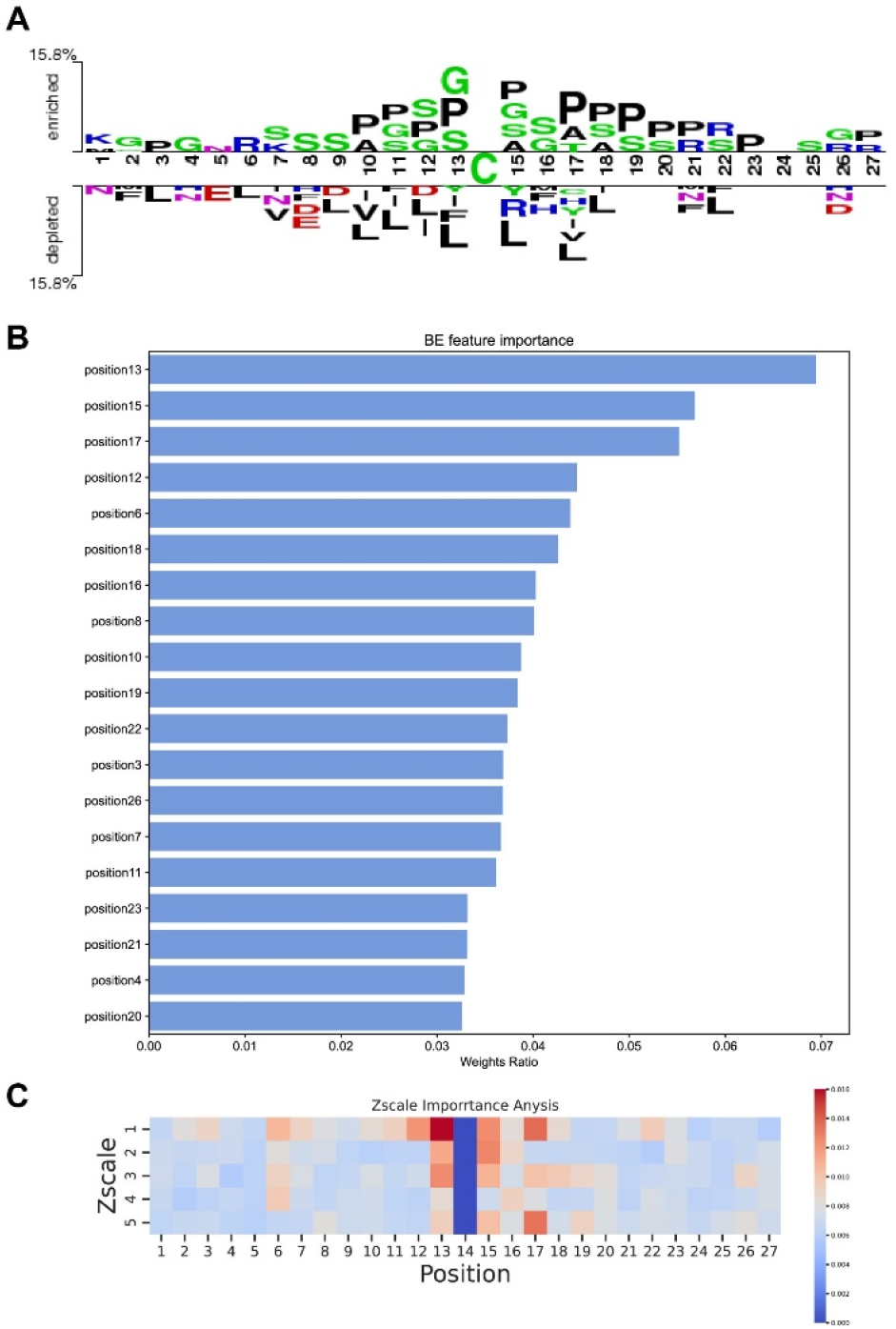
(A) Two sample logo of positive and negative data sets. (B) Feature importance of BE. (C) Feature importance of ZSCALE

### 3.4 Construction of SBP-SITA Web-server

We developed an easy-to-use online tool for the prediction of the S-itaconation sites, dubbed Sequence-based predictor of S-itaconation (SBP-SITA). It contains the Ensemble model in this article, and accepts input protein sequences as the FASTA format directly or uploads the sequence file (Figure 4). After the job submission, the prediction will start and take several minutes. Finally, the prediction results are output in tabular form with five columns: protein, position, sequence, prediction score.

## 4 Conclusion and discussion

S-itaconation is a significant biological activity in the activation of macrophage cells, regulating many pathways. Recently, thousands of serine S-itaconation sites have been found in mouse macrophages (Qin *et al*., 2019). Our study constructed the first S-itaconation prediction tools named SBP-SITA. It was an Ensemble classifier integrated six machine learning models, achieving 0.73 AUC, and could be accessible at the website (https://bioinfogo.org/sbp-sita/).

In feature exploration, the Z1 of the Zscale is more significant to others (Figure 3C), it reflects the polarity of amino acids, but P is not a polar amino acid, so the sterically hindered of amino acid side chains may influence the occurrence of S-itaconation. But the detailed mechanism needs to be further explored.

## 7 Supplementary Material

**Table S 1.** Z-scale value

| Amino acid | Z1 | Z2 | Z3 | Z4 | Z5 | Amino Acid | Z1 | Z2 | Z3 | Z4 | Z5 |
| --- | --- | --- | --- | --- | --- | --- | --- | --- | --- | --- | --- |
| A | 0.24 | -2.32 | 0.60 | -0.14 | 1.30 | M | -2.85 | -0.22 | 0.47 | 1.94 | -0.98 |
| C | 0.84 | -1.67 | 3.71 | 0.18 | -2.65 | N | 3.05 | 1.60 | 1.04 | -1.15 | 1.61 |
| D | 3.98 | 0.93 | 1.93 | -2.46 | 0.75 | P | -1.66 | 0.27 | 1.84 | 0.70 | 2.00 |
| E | 3.11 | 0.26 | -0.11 | -3.04 | -0.25 | Q | 1.75 | 0.50 | -1.44 | -1.34 | 0.66 |
| F | -4.22 | 1.94 | 1.06 | 0.54 | -0.62 | R | 3.52 | 2.50 | -3.50 | 1.99 | -0.17 |
| G | 2.05 | 4.06 | 0.36 | -0.82 | -0.38 | S | 2.39 | -1.07 | 1.15 | -1.39 | 0.67 |
| H | 2.47 | 1.95 | 0.26 | 3.90 | 0.09 | T | 0.75 | -2.18 | -1.12 | -1.46 | -0.40 |
| I | -3.89 | -1.73 | -1.71 | -0.84 | 0.26 | V | -2.59 | -2.64 | -1.54 | -0.85 | -0.02 |
| K | 2.29 | 0.89 | -2.49 | 1.49 | 0.31 | W | -4.36 | 3.94 | 0.59 | 3.44 | -1.59 |
| L | -4.28 | -1.30 | -1.49 | -0.72 | 0.84 | Y | -2.54 | 2.44 | 0.43 | 0.04 | -1.47 |

## Reference

Chen, Z., He, N., Huang, Y., Qin, W.T., Liu, X., and Li, L. (2018a). Integration of A Deep Learning Classifier with A Random Forest Approach for Predicting Malonylation Sites. Genomics, proteomics & bioinformatics 16, 451–459. 10.1016/j.gpb.2018.08.004.

Chen, Z., Liu, X., Li, F., Li, C., Marquez-Lago, T., Leier, A., Akutsu, T., Webb, G.I., Xu, D., Smith, A.I., et al. (2019). Large-scale comparative assessment of computational predictors for lysine post-translational modification sites. Briefings in bioinformatics 20, 2267–2290. 10.1093/bib/bby089.

Chen, Z., Zhao, P., Li, F., Leier, A., Marquez-Lago, T.T., Wang, Y., Webb, G.I., Smith, A.I., Daly, R.J., Chou, K.C., and Song, J. (2018b). iFeature: a Python package and web server for features extraction and selection from protein and peptide sequences. Bioinformatics 34, 2499–2502. 10.1093/bioinformatics/bty140.

Chen, Z., Zhao, P., Li, F., Marquez-Lago, T.T., Leier, A., Revote, J., Zhu, Y., Powell, D.R., Akutsu, T., Webb, G.I., et al. (2020). iLearn: an integrated platform and meta-learner for feature engineering, machine-learning analysis and modeling of DNA, RNA and protein sequence data. Briefings in bioinformatics 21, 1047–1057. 10.1093/bib/bbz041.

Friedman, J.H. (2001). Greedy function approximation: A gradient boosting machine. The Annals of Statistics 29, 1189–1232, 1144.

Huang, Y., He, N., Chen, Y., Chen, Z., and Li, L. (2018). BERMP: a cross-species classifier for predicting m(6)A sites by integrating a deep learning algorithm and a random forest approach. International journal of biological sciences 14, 1669–1677. 10.7150/ijbs.27819.

Lyu, X., Li, S., Jiang, C., He, N., Chen, Z., Zou, Y., and Li, L. (2020). DeepCSO: A Deep-Learning Network Approach to Predicting Cysteine S-Sulphenylation Sites. Front Cell Dev Biol 8, 594587. 10.3389/fcell.2020.594587.

Mills, E.L., Ryan, D.G., Prag, H.A., Dikovskaya, D., Menon, D., Zaslona, Z., Jedrychowski, M.P., Costa, A.S.H., Higgins, M., Hams, E., et al. (2018). Itaconate is an anti-inflammatory metabolite that activates Nrf2 via alkylation of KEAP1. Nature 556, 113–117. 10.1038/nature25986.

Ning, W., Jiang, P., Guo, Y., Wang, C., Tan, X., Zhang, W., Peng, D., and Xue, Y. (2020). GPS-Palm: a deep learning-based graphic presentation system for the prediction of S-palmitoylation sites in proteins. Briefings in bioinformatics. 10.1093/bib/bbaa038.

O’Neill, L.A.J., and Artyomov, M.N. (2019). Itaconate: the poster child of metabolic reprogramming in macrophage function. Nature reviews. Immunology 19, 273–281. 10.1038/s41577-019-0128-5.

Qin, W., Qin, K., Zhang, Y., Jia, W., Chen, Y., Cheng, B., Peng, L., Chen, N., Liu, Y., Zhou, W., et al. (2019). S-glycosylation-based cysteine profiling reveals regulation of glycolysis by itaconate. Nature chemical biology 15, 983–991. 10.1038/s41589-019-0323-5.

Qin, W., Zhang, Y., Tang, H., Liu, D., Chen, Y., Liu, Y., and Wang, C. (2020). Chemoproteomic Profiling of Itaconation by Bioorthogonal Probes in Inflammatory Macrophages. Journal of the American Chemical Society 142, 10894–10898. 10.1021/jacs.9b11962.

Sandberg, M., Eriksson, L., Jonsson, J., Sjostrom, M., and Wold, S. (1998). New chemical descriptors relevant for the design of biologically active peptides. A multivariate characterization of 87 amino acids. Journal of medicinal chemistry 41, 2481–2491. 10.1021/jm9700575.

Strelko, C.L., Lu, W., Dufort, F.J., Seyfried, T.N., Chiles, T.C., Rabinowitz, J.D., and Roberts, M.F. (2011). Itaconic acid is a mammalian metabolite induced during macrophage activation. Journal of the American Chemical Society 133, 16386–16389. 10.1021/ja2070889.

Tasci, E., Uluturk, C., and Ugur, A. (2021). A voting-based ensemble deep learning method focusing on image augmentation and preprocessing variations for tuberculosis detection. Neural Comput Appl, 1–15. 10.1007/s00521-021-06177-2.

Vacic, V., Iakoucheva, L.M., and Radivojac, P. (2006). Two Sample Logo: a graphical representation of the differences between two sets of sequence alignments. Bioinformatics 22, 1536–1537. 10.1093/bioinformatics/btl151.

Wang, H., Wang, Z., Li, Z., and Lee, T.Y. (2020). Incorporating Deep Learning With Word Embedding to Identify Plant Ubiquitylation Sites. Front Cell Dev Biol 8, 572195. 10.3389/fcell.2020.572195.

Weiss, J.M., Davies, L.C., Karwan, M., Ileva, L., Ozaki, M.K., Cheng, R.Y., Ridnour, L.A., Annunziata, C.M., Wink, D.A., and McVicar, D.W. (2018). Itaconic acid mediates crosstalk between macrophage metabolism and peritoneal tumors. The Journal of clinical investigation 128, 3794–3805. 10.1172/JCI99169.

Weng, S.L., Kao, H.J., Huang, C.H., and Lee, T.Y. (2017). MDD-Palm: Identification of protein S-palmitoylation sites with substrate motifs based on maximal dependence decomposition. PloS one 12, e0179529. 10.1371/journal.pone.0179529.

